# Love, not food, could have paved the path for dog domestication: A lesson from free-ranging dogs

**DOI:** 10.1101/161596

**Authors:** Debottam Bhattacharjee, Shubhra Sau, Jayjit Das, Anindita Bhadra

## Abstract

Dogs (*Canis lupus familiaris*) are the first species to have been domesticated, and unlike other domesticated species, they have developed a special bonding with their owners. The ability to respond to human gestures and language is a key factor in the socio-cognitive abilities of dogs that have made them our best friend. Free-ranging dogs provide an excellent model system for understanding the dog domestication process. In India, free-ranging dogs occupy every possible human habitation, and interact with humans regularly. They scavenge among garbage, beg for food from humans, give birth in dens close to human habitations, and establish social bonds with people. However, there is ample dog-human conflict on the streets, leading to morbidity and mortality. Hence the ability to assess an unfamiliar human before establishing physical contact could be adaptive for dogs especially in the urban environment. We tested a total of 103 adult free-ranging dogs to investigate their response to immediate and long-term food and social rewards. The dogs were provided a choice of obtaining a food reward either from the hand or the ground. The dogs avoided making physical contact with the unfamiliar human. While immediate rewards were not effective in changing this response, the long-term test showed a strong effect of the social reward on the response of dogs. Our results revealed that dogs tend to build trust based on affection, and not food rewards. This study provides significant insights into nuances of the dynamics that could have paved the path to dog domestication.

## Introduction

Living in close proximity with humans can have several adaptive advantages for animals, while posing challenges for survival at the same time. Human habitations can be good sources for food, shelter and protection; and many species of animals, from insects to mammals are known to have adapted to co-habiting with humans, as pests, parasites, commensals and domesticates (Castillo et al., 2003; Pocock et al., 2004; Vannier-Santos and Lenzi, 2011). The changing landscape of human habitation, from more rural to more urban over the past decades has led to an increasing interest in urban ecosystems (Alberti et al., 2008; Mcintyre and Hope, 2008; Pickett et al., 2008) Species that adapt to the urban environment present interesting case studies to understand how they resolve issues of conflict with humans while exploiting new niches created in these human dominated landscapes. This question is all the more interesting for species like birds and mammals, in which decision making might be influenced by their experiences from interactions with humans in the urban space (Ditchkoff et al., 2006; Maklakov et al., 2011; White et al., 2005).

Most of the species present in urban areas are generalists, omnivorous and have higher tolerance to human disturbance (Grimm et al., 2008; Lizee et al., 2011; Shochat et al., 2006). Successful urban species typically show plasticity in behaviours that help them to adjust to major environmental disturbances and exploit new niches. Yet, most urban species maintain a wary distance with humans, showing flight response to human approach and close interactions (Carrete and Tella, 2011; Møller A, 2008; Rodewald and Shustack, 2008). Studies on several urban adapted mammals have shown that though they scavenge in human settlements, they prefer to den away from humans, avoiding human proximity during whelping (Ross et al., 2010; Theuerkauf and Jedrzejewski, 2002; Ye et al., 2007). The dog (*Canis lupus familiaris*) has shared space and coevolved with humans for centuries, being the first ever species to be domesticated by humans (Clutton-Brock, 1995). Though the domestic dog is mostly recognized as pets, free-ranging dogs comprise almost 80% of the world’s dog population (Boitani and Ciucci, 1995; Hughes and Macdonald, 2013), and are an integral part of the human environment in most developing countries (Vanak and Gompper, 2009). They experience differing levels of human interactions, both positive and negative (Bhattacharjee et al, *In Press*), and are largely dependent on human generated wastes for their sustenance (Majumder et al., 2014). Free-ranging dogs are not under direct human supervision (Cafazzo et al., 2010; Majumder et al., 2014; Vanak and Gompper, 2009), but have adapted to living in close proximity with humans as commensals. They are also considered to be reservoirs of various zoonotic diseases including rabies and hence a threat to both humans and wildlife (Butler et al., 2004; Fekadu, 1982). Moreover, they scatter garbage, defecate in open spaces and disturb people by their nocturnal barking. Hence they are considered to be a nuisance by a part of the human population, though they are often cared for by some. It is likely that their experiences of interactions with humans influence their behaviour to some extent, determining their perception of humans in general.

In India, free-ranging dogs have existed as a continuous population for centuries (Thapar R, 1990). They have a ubiquitous presence from remote villages to metropolitan areas. They are primarily scavengers, but are known to hunt in packs in the fringes of human habitations too (Kumar and Paliwal, 2015; Young et al., 2011). They live in stable social packs that show interesting cooperation-conflict dynamics, especially over pup rearing (Majumder et al., 2014; Paul and Bhadra, 2017; Paul et al., 2014; Paul et al., 2015). Early life mortality is very high in spite of extensive parental and alloparental care, with only 19% of the pups reaching adulthood (Paul et al., 2016). 63% of this mortality is human induced, and yet, the free-ranging dogs do not avoid human proximity during whelping, but often use spaces within human habitations as dens (Sen Majumder et al., 2016). They show plasticity in their interactions with humans; pups readily follow human pointing gestures, but juveniles tend not to do so. Adults use reliability cues to adjust their responses to human pointing and are good at retrieving food from human artefacts like closed garbage bags (Bhattacharjee et al, *In press*). Humans are thus a source of food and shelter but also responsible for mortality of the free-ranging dogs, and thus it is imperative for the dogs to assess the intentions of humans before interacting with them. It has been suggested that canines have a predisposition to attend to the actions of their social companions, provided they must learn to recognize humans as companions and understand the relationship through learning and experience (Reid, 2009). Moreover, social attachment with humans is considered to have been a key factor in dog domestication (Nagasawa et al., 2015). Thus, understanding the ability of the free-ranging dogs to establish a social connection with unfamiliar humans can help to shed light on the domestication process. Here we investigate the dog – human relationship in the context of food and social rewards in urban environment.

A study by Feuerbacher and Wynne (2012) concluded no effect of a brief social reward on pet and shelter dogs as compared to food reward (Feuerbacher and Wynne, 2012), but contextual differences might play a determining role in case of the free-ranging dogs. Another study showed pet dogs’ tendency to prefer food to petting; but petting seemed to be important when it was compared with vocal praise (Feuerbacher and Wynne, 2014). Hence, without considering the effect of different environmental conditions and life experiences, direct comparison of outcomes from pet dogs with the free-ranging dogs would not be valid. We conducted field trials on free-ranging dogs to test the effect of food and social rewards on their tendency to make contact with unfamiliar humans. Our experiments comprised of both one-off trials with brief exposure of social petting, as well as long-term repeated trials for both kinds of rewards. We provided dogs with a choice to obtain food from either a human hand or the ground. Since free-ranging dogs are scavengers, and they also receive negative interactions from humans, we hypothesized that they would prefer to take food from the ground, rather from the experimenter’ s hand. We expected that the immediate social reward would increase the dogs’ tendency to take food from the hand. However, since pet dogs respond more to food than to petting, we expected the free-ranging dogs to show an increased tendency to feed from the hand on being provided with long term food, rather than social rewards.

## Materials and methods

### Subjects and study area

We tested a total of 103 adult free-ranging dogs located randomly in different urban areas - Mohanpur (22°56’49” N and 88°32’4”E), Kalyani (22°58’30”N, 88°26’04”E)and Kolkata (22°57’26”N, 88°36’39”E), West Bengal, India. Sexes of the dogs were determined by looking at their genitals. All individuals were photographed for record and tracking purpose. The individuals were tracked for the long term experiments using their location and morphological features like coat colour, patch patterns and any other distinguishing features in the body.

### Experimental procedure

#### (i)One off Test

We used 30 random adult individual dogs to test their tendency to approach unfamiliar humans for food. A single piece of raw chicken weighing approximately 10-15 gm was used as the food reward. In trial 1, the experimenter (E) placed a food reward on the palm of his hand (hand chosen randomly) and held it open close to the ground, at a height of 5-10 cm. He placed another similar piece of raw chicken at the same time, but on the ground in front of him, such that there was a distance of 0.6 m between the two reward options. The set-up was designed such that both the reward options were equally accessible to the dog (Supplementary Information 1: Fig S1, Video S1). E tried to attract the attention of an individual dog using sounds that are typically used to call out to dogs on streets in India (Supplementary Information 2: Video S2) for 1-2 sec, from an approximate distance of 4 ft. Since these dogs were not on leash, we tried to ensure that the distance remained roughly the same before recording the trials. The response of the dog was video recorded for 1 min or till an individual made a choice, whichever was earlier. E kept gazing at the dog throughout the trial, and the same person carried out all the trials. After completion of trial 1, E provided the dog with a social reward by petting 3 times on its head. After an interval of 5-10 seconds, trial 2 was run, where the individuals again had to make a choice from the same set up as in trial 1.

The control trials were exactly the same as the test trials but here, E did not provide any social reward in between trials 1 and 2. A separate set of 30 adult dogs were tested in this condition.

#### (ii)Long term test

A total of 43 adult dogs different from the one-off test and control conditions were randomly selected from diverse locations for a long term experiment in order to investigate the effect of learning in the context of food and social rewards. From this set of 43 individuals, two subsets were randomly generated; 21 individuals for food and 22 individuals for social rewards. Each dog was tested a total of six times, at increasing intervals of 1, 2, 3, 4 and 5 days. Thus, for every dog, the experiment commenced on Day 0, and was conducted on Days 1, 3, 6, 10 and 15 respectively. Unlike the previous two conditions, in this case, only trial 1 was run, keeping the protocol constant. Additional food (one piece of chicken) or social (petting thrice on the head) rewards were provided by E, 1 min before the trial, to the respective subsets of dogs except on Day 0. On Day 0, no additional food or social reward was provided, such that this represented the response of the naïve dogs to E.

### Data analysis

All the videos were coded by a single experimenter different from E, and the data was used for further analysis. We used Shapiro-Wilk tests to check for normality of our data and found them to be not normally distributed, thus we performed non-parametric tests.

We considered all the “naïve” responses - trial 1 of the one off-test and control conditions and the Day 0 responses of long term observations in order to investigate the population level tendency of free-ranging dogs to make attachment with humans for food. We compared the number of dogs that obtained food from the ground and human hand using Goodness of Fit Chi-square test. We defined latency as the time interval between catching the attention of an individual and its approach to either of the options provided. We compared latencies of the dogs that obtained food from the ground and from the hand using Mann Whitney U test. We identified the sexes of the dogs and compared the response of the two sexes in obtaining food rewards using Contingency chi-square test. We calculated all possible combinations (hand-hand, ground-hand, ground-ground and hand-ground) of obtaining food reward by dogs between trial 1 and 2 of both test and control conditions. Hand – hand and ground – ground situations were considered as “no change” and hand – ground and ground – hand situations as “change”. We then compared change and no change categories for both test and control trials by using goodness of fit chi-square tests. We compared latencies of dogs between trial 1 and 2 for both the test and control conditions using Wilcoxon paired-sample tests.

We built a socialization index based on the vigour of tail wagging and gazing at different food reward options by the dogs (Table 1). Tail wagging by free-ranging dogs establish an affirmative association with humans and indicative of positive social bond. Gazing or alternation of gaze indicate dogs’ hesitant nature to approach any of the food options. Scores were assigned in such a way that an individual could have a maximum index value of 8 by showing rapid tail wagging and no gazing at all.

**Table 1.** Socialization index incorporating the behaviours shown by dogs and corresponding scores. Socialization index was built based on tail wagging and gazing behaviour. Definitions of different types of tail wagging and gazing behaviours and their corresponding scores within a range of 1 to 4 provided.

| Tail Wagging |  | Gazing |  |
| --- | --- | --- | --- |
| Type | Score | Type | Score |
| Rapid (tail wagging along with back movement) | 4 | No | 4 |
| Fast (without back movement) | 3 | At hand food | 3 |
| Slow | 2 | At both hand and ground food | 2 |
| No | 1 | At ground food | 1 |

We compared the index values of individuals across the two trials for both test and control conditions using Wilcoxon paired-sample tests. Generalized linear mixed models (GLMM) with binomial distributions were used to check any effect of day intervals, latencies, and socialization index values over dogs’ preference for obtaining food reward from either hand or ground in long-term experiments.

Separate models (GLMM) were built for the two different conditions where additional food and social rewards were provided. Identity of individuals was included as a random effect on the intercept. We used AIC values for comparison in order to get the best-fitting models. We determined consistency of individuals obtaining food reward from hand and ground at 100% and 80% levels for both the long-term experiments. Similarly, overall inconsistency was calculated where dogs changed their preference on every alternate day interval.

A second coder naïve to the purpose of the study coded 20% of the data to check inter-rater reliability. It was perfect food preference (cohen’s kappa = 1.00) and socialization index building (cohen’s kappa = 1.00) and almost perfect for latency (cohen’s kappa = 0.95). The alpha level was 0.05 throughout the analysis. GLMMs were performed using “lme4” package of R Studio (R Development Core Team, 2015). Along with R, other statistical analyses were performed using StatistiXL version 1.11.0.0.

## Results

### (i)Attachment of dogs to humans

Considering the “naïve” responses of all 103 individuals, 37% obtained food reward from the human hand and 63% from the ground, thereby showing a bias against making physical contact with E (Goodness of fit; χ^2^ = 7.078, df = 1, p = 0.008, Fig 1).

**Fig 1.**
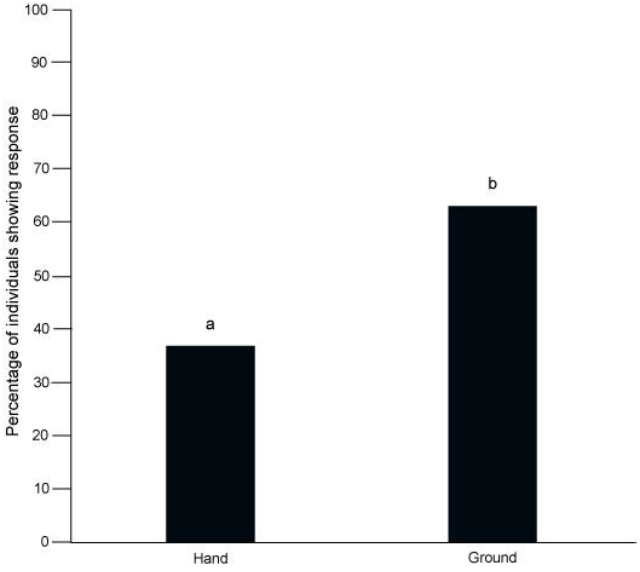
Bar graph showing percentage of individuals that obtained food reward from human hand and ground out of all naïve responses. Dogs showed a significantly higher tendency to obtain the reward from the ground (Goodness of fit; χ^2^ = 7.078, p = 0.008). “a” and “b” indicate significant differences between the categories.

However, we did not see any difference in latencies to approach between the responders who showed different choices (Mann Whitney U test; U = 1301.00, df1 = 38, df2 = 65, p = 0.65), which suggests that the final choice did not influence the time taken to reach a decision to respond. The two sexes were comparable in their preference to obtain food from either the hand or the ground (Contingency χ^2^; χ^2^ = 1.573, df = 1, p = 0.21).

### Effect of immediate social reward/petting

We found that in the one-off test, 73% and 83% of the individuals showed “no change” in their response in trial 2 for the test and control conditions respectively. For both the conditions, proportion of “no change” responses were significantly higher than “change” (Test - Goodness of fit; χ^2^ = 6.533, df = 1, p = 0.01: Control - χ^2^ = 13.333, df = 1, p < 0.001), suggesting no effect of immediate social reward (Fig 2).

**Fig 2.**
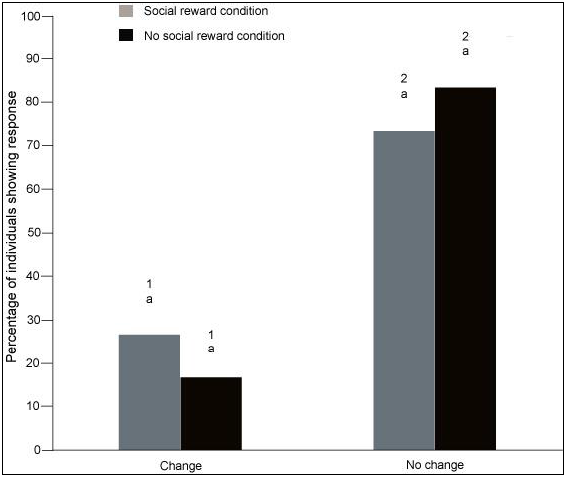
Bar graph illustrating percentage of individuals showing “change” and “no change” responses for preference of food reward in the 2^nd^ trials of test and control conditions. “No change” indicates same preference of obtaining food in trials 1 and 2 (Hand – Hand, Ground - Ground). Change indicates a change or switch in preference from trial 1 to trial 2 (Hand – Ground, Ground - Hand). Grey bars indicate responses in the test (social reward) condition and black bars indicate responses in the control (no social reward) condition. Letter “a” indicates no significant differences within the categories (within change and within no change). “1” and “2” indicate significant differences between the categories (between change and no change).

Interestingly, we found faster approach by dogs to the set-up in trial 2 for the test condition; thus, the latency significantly decreased when social reward was provided (Wilcoxon paired-sample test; T = 86.00, N = 30, p = 0.004), but it remained unchanged in the control condition, when no social reward was provided (Wilcoxon paired-sample test; T = 166.500, N = 30, p = 0.427, Fig 3). Since separate sets of individuals were present for the test and control conditions, we compared the latencies of 1st trials and 2^nd^ trials between test and control conditions. We found no difference for the 1^st^ trial latencies of the two conditions (Mann Whitney U test; U = 480.00, df = 30, df = 30, p = 0.66), but noticed a significant difference between the 2^nd^ trial latencies (Mann Whitney U test; U = 584.500, df = 30, df = 30, p = 0.04). These results suggest that the dogs might be prone to showing a stronger response to humans when they receive positive social interactions. However, we did not find any difference in socialization index values between 1^st^ and 2^nd^ trials for both test (Wilcoxon paired-sample test; T = 73.500, N = 30, p = 0.14) and control conditions (Wilcoxon paired-sample test; T = 111.500, N = 30, p = 0.20).

**Fig 3.**
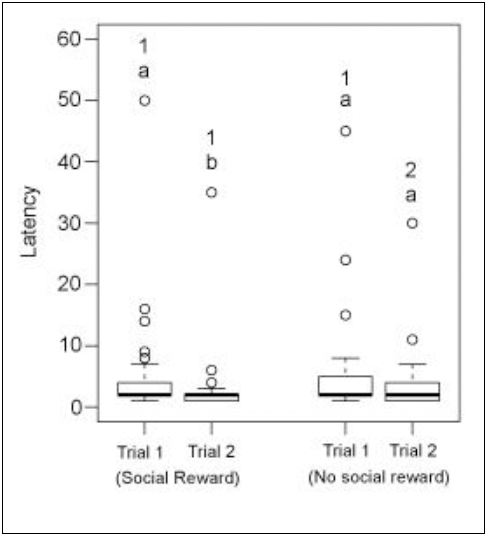
Box and whiskers plot showing the latency to approach the set-up. Dogs showed significant difference in latency between 1^st^ and 2^nd^ trials for the test (social reward) condition. Latency remained comparable between 1^st^ and 2^nd^ trials in the control (no social reward) condition. Boxes represent interquartile range, horizontal bars within boxes indicate median values, and whiskers represent the upper range of the data. “a” and “b” indicate significant differences within the categories (within social reward and within no social reward). “1” and “2” indicate significant differences between the categories (between social and no social reward).

### Effect of long-term social and additional food rewards

Based on AIC values, the best-fitting model depicted socialization index to be the only significant predictor for dogs’ preference of obtaining food in case of the long-term experiment with additional food reward (Fig 4, Supplementary Information 3). On the other hand, we found socialization index, latency and time intervals as significant predictors affecting the response for the long-term experiment with social rewards (Fig 5, Supplementary Information 4). In the long-term additional food reward experiment, 11 out of 21 individuals (52%) were 100% consistent for obtaining food reward from the ground, whereas only a single individual consistently obtained food from the hand (Goodness of fit; χ^2^ = 8.333, df = 1, p = 0.004, Fig 6). In the long-term social reward experiment, 3 out of 22 (14%) and 8 out of 22 (36%) individuals showed 100% consistency at obtaining food from ground and hand respectively (Goodness of fit; χ^2^ = 2.273, df = 1, p = 0.13, Fig 7). We found a difference in the 100% consistency levels of obtaining food from hand and ground between social and additional food reward conditions (Contingency chi-square; χ^2^ = 9.991, df = 1, p = 0.002). At 80% consistency level, 3 out of 21 (14%) and 1 out of 21 (5%) individuals obtained food reward from hand and ground (Goodness of fit; χ^2^ = 1.000, df = 1, p = 0.32) respectively for additional food reward condition; whereas, none of the individuals from social reward condition showed 80% consistency. 1 out of 21 individuals for additional food and 1 out of 22 individuals for social reward condition changed their responses on every alternate day of the experiment, thereby showing inconsistency.

**Fig 4.**
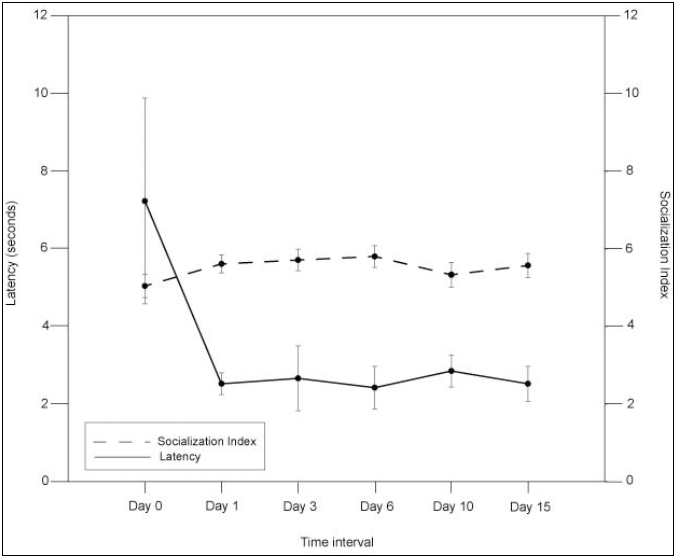
Average latency and socialization index values (± SE) at different intervals (in days) for long-term additional food reward condition. The primary y-axis depicts latency and secondary y-axis depicts socialization index with x-axis showing the specific time interval. The solid line indicates latency and the dashed line indicates socialization index.

**Fig 5.**
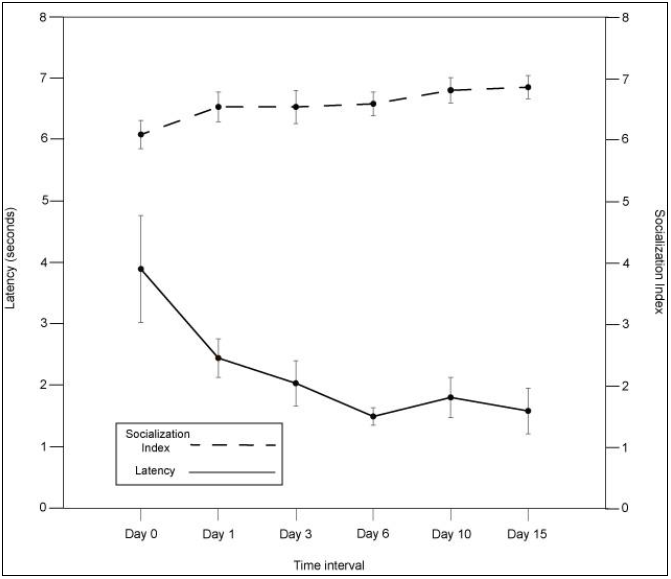
Average latency and socialization index values (± SE) at the day intervals for long-term social reward condition. The primary y-axis depicts latency and secondary y-axis depicts socialization index with the x-axis showing the specific time interval. The solid line indicates latency and dashed line indicates socialization index.

**Fig 6.**
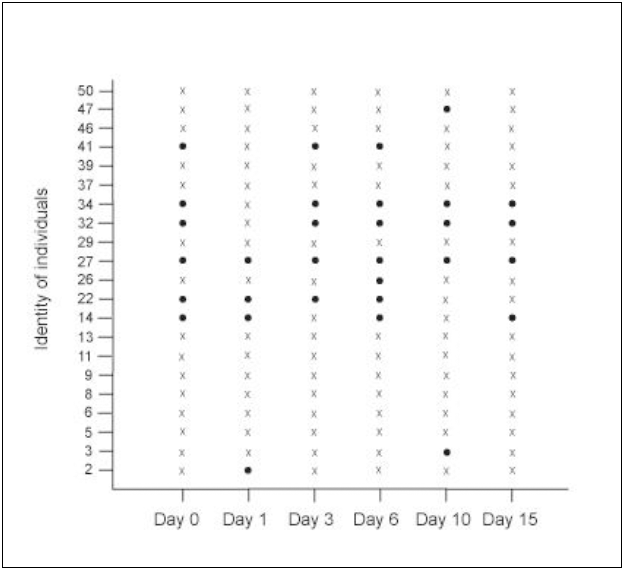
Identity of individuals and their preference of obtaining reward at specific day intervals for the long-term additional food reward condition. The solid dots indicate obtaining food from hand and the crosses indicate obtaining food from ground. Day 0 responses are naïve as dogs had no previous exposure to additional food reward.

**Fig 7.**
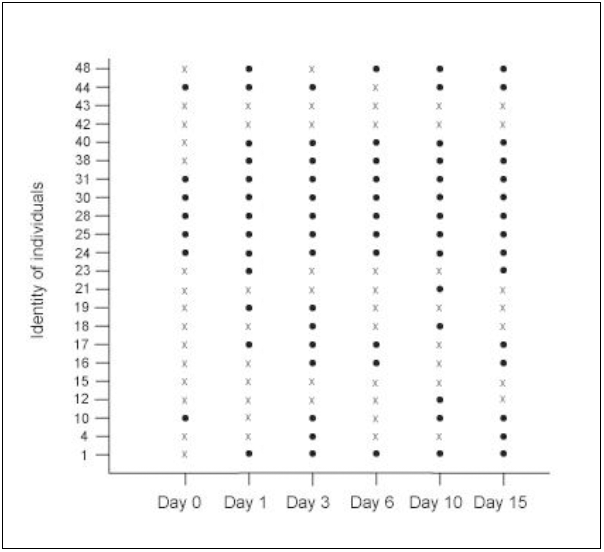
Identity of individuals and their preference of obtaining reward at specific day intervals for the long-term social reward condition. The solid dots indicate obtaining food from hand and the crosses indicate obtaining food from ground. Day 0 responses are naïve as dogs had no previous exposure to social reward.

## Discussion

Free-ranging dogs demonstrated a bias against making physical contact with unfamiliar humans, as suggested by the higher proportion of individuals opting to take the food from the ground, validating our hypothesis. In the one-off test, dogs elicited a significantly faster response (reduced latency) or an increased tendency to approach the set-up when the social reward was provided. However, contrary to our expectations, a single positive feedback in terms of social reward did not prove to be substantial in establishing trust in the unfamiliar human. Moreover, it did not translate into an increased tendency to interact with the experimenter as there was no change in the socialization index between consecutive trials when the social reward was provided. The one-off experiments thereby reinforce the idea that free-ranging dogs are generally wary of humans (Bhattacharjee et al, *In press*), and prefer not to make physical contact with unfamiliar humans, even after receiving brief positive reinforcement through food or social rewards.

The long-term experiments provided an interesting insight into the free-ranging dogs’ relationship with humans, which was very different from our hypothesis based on the results obtained with pet dogs in the past(Feuerbacher and Wynne, 2012),^38^. Long-term provisioning of additional food reward increased the socialization index values of dogs who obtained food reward from the human hand. However, there was no significant reduction in the latency to respond, which suggests that the dogs were hesitant to make direct contact with humans, despite increasing familiarity and positive reinforcement with additional food reward. The most striking observation was the change in the dogs’ response to the experimenter in the presence of the long-term social reward. Dogs exposed to the social reward showed reduced latency (thus increased interest) to approach the experimenter and an increase in the socialization index. Moreover, the dogs’ preference to feed from the human hand increased with increased exposure to the social reward. The high degree of consistency shown by the dogs in obtaining food from the ground in the additional food reward condition further validates their predisposition to avoid physical contact with unfamiliar humans. In contrast, higher number of dogs showed 100% consistency in preferentially taking food from the human hand in the social reward condition. To summarize,long-term social reward, but not food reward, impacted the dogs’ tendency to make physical contact with humans, which suggests that social reward is more effective in building trust between dogs and unfamiliar humans than food rewards.

A recent study concluded that domestication has been a key factor contributing to dogs’ ability of visual and physical contact with humans (Nagasawa et al., 2015). It has been suggested that short-term sensory interactions between pet dogs and their owners influence hormonal levels (e.g. oxytocin) and heart rate (Handlin et al., 2011). Rising levels of oxytocin in dogs help to maintain their social orientation, affiliation and gazing toward their owners (Nagasawa et al., 2015; Romero et al., 2014), and gazing towards dogs increases the oxytocin levels in their owners, which in turn induces a rise in oxytocin in the dogs, thus strengthening the dog-human bonding (Nagasawa et al., 2015). Though free-ranging dogs tend to avoid direct human contact, studies have shown that they gaze and even seek help from strangers when faced with an unfamiliar task (Bhattacharjee et al., 2017). Free-ranging dogs are also known to regularly beg from humans, using the gazing behaviour (Bhadra and Bhadra, 2014). They often experience negative interactions with humans, and their tendency to avoid direct physical contact with unfamiliar humans could be a possible outcome of cumulative negative experiences. In our experiment with the long-term additional reward conditions, the dogs preferred to avoid contact when the experimenter provided additional food reward. However, long term social reward increased the dogs’ tendency to make physical contact with the experimenter. In their day-to-day lives, the free-ranging dogs routinely encounter unfamiliar humans, and the ability to assess human intentions could be highly adaptive under such circumstances. They are often lured with food and then beaten up or even poisoned by people (Paul et al., 2016) and hence, relying on strangers who offer food might have negative consequences for dogs. On the other hand, people who show affection to the dogs are less likely to harm them, and relying on such humans might indeed be advantageous.

In an earlier study, we have observed pups to readily follow human pointing, while juveniles and adults fail to do so. Interestingly, adults adjust their reliability on the experimenter based on immediate experience, choosing to follow pointing on positive reinforcement and refusing to follow pointing on negative reinforcement (Bhattacharjee et al. *In Press*). Thus, the free-ranging dogs show considerable plasticity in their tendency to follow human pointing, learning from experience through their development. The ability to follow human pointing is considered to be an important socio-cognitive ability in dogs, and has been suggested to be closely associated with their domestication (Hare and Tomasello, 2005; Miklósi et al., 2004; Miklösi et al., 1998; Paul et al., 2016; Soproni et al., 2001). The free-ranging dogs in India live very closely with humans, in every possible human habitation, and are dependent on humans for their sustenance, either directly or indirectly (Majumder et al., 2014; Sen Majumder et al., 2016; Vanak and Gompper, 2009). In contrast to other urban adapted animals, the free-ranging dogs have been observed to preferentially den close to humans, often within human homes (Sen Majumder et al., 2016). They interact with familiar as well as unfamiliar humans on a regular basis and receive both positive and negative interactions from them. During the early stages of domestication, ancestors of the present-day dogs would have been faced with a similar situation. Human communities would have acted as a lucrative source of nutrition, but the adventurous individuals which ventured too close could have been easy prey for the humans. Someone who threw a bone to a dog could have turned into a hunter in no time, but a human who put out a friendly hand to pet a dog would have been less likely to attack it. The oxytocin feedback that is known to help establish bonding between dogs and their owners could have been influential in this trust building phase of the relationship between the two species. We speculate that the tendency of dogs to rely more on positive social interactions, rather than food as a cue for trust building with unfamiliar humans could have acted as an important behavioural paradigm in the evolution of the dog-human relationship.

## Acknowledgements

The authors thank Mr. Susnata Karmakar, IISER Kolkata, for lending his vehicle for use during the field experiments.

## Author contributions

DB, SS, JD carried out the field experiments, SS was the constant Experimenter throughout the work. DB coded the videos and analyzed the data. AB and DB designed the protocol. AB supervised the work and co-wrote the paper with DB.

